# Novel color vision assessment tool: AIM Color Detection and Discrimination

**DOI:** 10.1101/2024.09.26.615300

**Authors:** Jingyi He, Jan Skerswetat, Peter J. Bex

## Abstract

Color vision assessment is essential in clinical practice, yet different tests exhibit distinct strengths and limitations. Here we apply a psychophysical paradigm, Angular Indication Measurement (AIM) for color detection and discrimination. AIM is designed to address some of the shortcomings of existing tests, such as prolonged testing time, limited accuracy and sensitivity, and the necessity for clinician oversight. AIM presents adaptively generated charts, each a N×M (here 4×4) grid of stimuli, and participants are instructed to indicate either the orientation of the gap in a cone-isolating Landolt C optotype or the orientation of the edge between two colors in an equiluminant color space. The contrasts or color differences of the stimuli are adaptively selected for each chart based on performance of prior AIM charts. In a group of 23 color-normal and 15 people with color vision deficiency (CVD), we validate AIM color against Hardy-Rand-Rittler (HRR), Farnsworth-Munsell 100 hue test (FM100), and anomaloscope color matching diagnosis and use machine learning techniques to classify the type and severity of CVD. The results show that AIM has classification accuracies comparable to that of the anomaloscope, and while HRR and FM100 are less accurate than AIM and an anomaloscope, HRR is very rapid. We conclude that AIM is a computer-based, self-administered, response-adaptive and rapid tool with high test-retest repeatability that has the potential to be suitable for both clinical and research applications.

## Introduction

Color vision deficiency (CVD), or colorblindness, is characterized by a reduced ability to detect and discriminate colors. This deficit can be categorized into two primary subtypes: inherited CVD and acquired CVD (Simunovic, 2016). In inherited CVD, genetic mutations can result in various types and degrees of severity (Deeb, 2004; Neitz & Neitz, 2000). As people with typical color vision are named trichromats, those with one defective cone type are referred to as anomalous trichromats, while those lacking one cone type are known as dichromats. Depending on the specific cone type affected— L (long-wavelength-sensitive)-, M (medium-wavelength-sensitive)-, or S (short-wavelength-sensitive)-cones— these conditions are respectively termed protan, deutan, and tritan (Bosten, 2019; Sharpe, Stockman, Jägle, & Nathans, 1999).

Color vision testing is a crucial component of eye examinations, aimed at screening for CVD. The reference standard for testing CVD is the anomaloscope, equipped with Rayleigh match (Thomas & Mollon, 2004) for screening red-green deficiencies and sometimes with Moreland match (Moreland, 2004) for screening blue-green deficiencies. Because of the efficiency in rapid assessments, pseudoisochromatic plates such as the Ishihara plates (Birch, 1997b) and the Hardy– Rand–Rittler (HRR) plates (Bailey, Neitz, Tait, & Neitz, 2004) are widely used for screening, however, they provide only a coarse classification and require an administrator to perform and monitor the test. Other color arrangement tests (Evans, Rodriguez-Carmona, & Barbur, 2021; Farnsworth, 1943) and computer-based tests with luminance noise added (Barbur, Harlow, & Plant, 1994; Mollon & Regan; Rabin, 1996), and children friendly tests (Tang et al., 2022) are more sensitive than pseudoisochromatic plates but take longer to complete. These tools each have distinct advantages and disadvantages that we recently discussed in He, Bex, & Skerswetat (2023). In brief, pseudoisochromatic plate tests are rapid but insensitive to mild CVD or change in CVD and provide only pass/fail classification, whereas psychophysical detection, matching and arrangement tests are more diagnostic and sensitive but impose a considerable time burden. Furthermore, all tests require trained personnel to administer and interpret results to assess vision loss.

Here we introduce AIM (Angular Indication Measurement) *Color Detection* and *Discrimination* tasks. AIM is a computer-based, rapid, self-administered paradigm (Neupane, Skerswetat, & Bex, 2024; Skerswetat, He, et al., 2024; Skerswetat, Ross, et al., 2024) that we adapt in this study for the assessment of color vision. The AIM paradigm is capable of measuring human visual performance and deriving the corresponding visual functions, including visual acuity (Skerswetat et al., 2024), stereopsis (Neupane et al., 2024), contrast sensitivity, motion and form coherence (Skerswetat, Ross, et al., 2024). This paradigm employs an adaptive procedure and adjusts visual stimuli based on the previous responses from the observer. The tasks require the observer to indicate the orientation or direction of stimuli (e.g., the “gap” in a Landolt C or the central line of a bipartite patch) around an outer response circle using an indication device such as a computer mouse. The angular error (i.e., angular difference between the indicated orientation and the actual orientation) provides an estimate of accuracy and precision as a function of stimulus intensity, and can also be converted to a forced-choice response by defining a corresponding error range, e.g. ±90° for 2-Alternative-Forced-Choice (AFC) or ±45° for 4-AFC etc.. Subsequently, a psychometric function can be derived based on a fit to the angular error as a function of the tested stimulus intensity. From this function, we calculate three key metrics: the threshold, the slope, and the minimum report error, which we termed ‘noise’, representing the sum of extrinsic and intrinsic noise sources (see Figure 1d), each providing insight into the observer’s visual processing and response capabilities (Skerswetat, He, et al., 2024). For the AIM Color Detection task, we use cone-isolating Landolt C stimuli for targeting standard L-, M-, and S-cone types (Stockman & Sharpe, 2000; Stockman, Sharpe, & Fach, 1999), respectively assessing detection thresholds to reveal the functionality of each cone type. For the AIM Color Discrimination task, we use a bipartite circle with different colors in each semicircle to measure color discrimination thresholds along eight color directions on an equiluminant color plane (Derrington, Krauskopf, & Lennie, 1984). Together, the two tasks provide a comprehensive assessment of color visual function.

**Figure 1.**
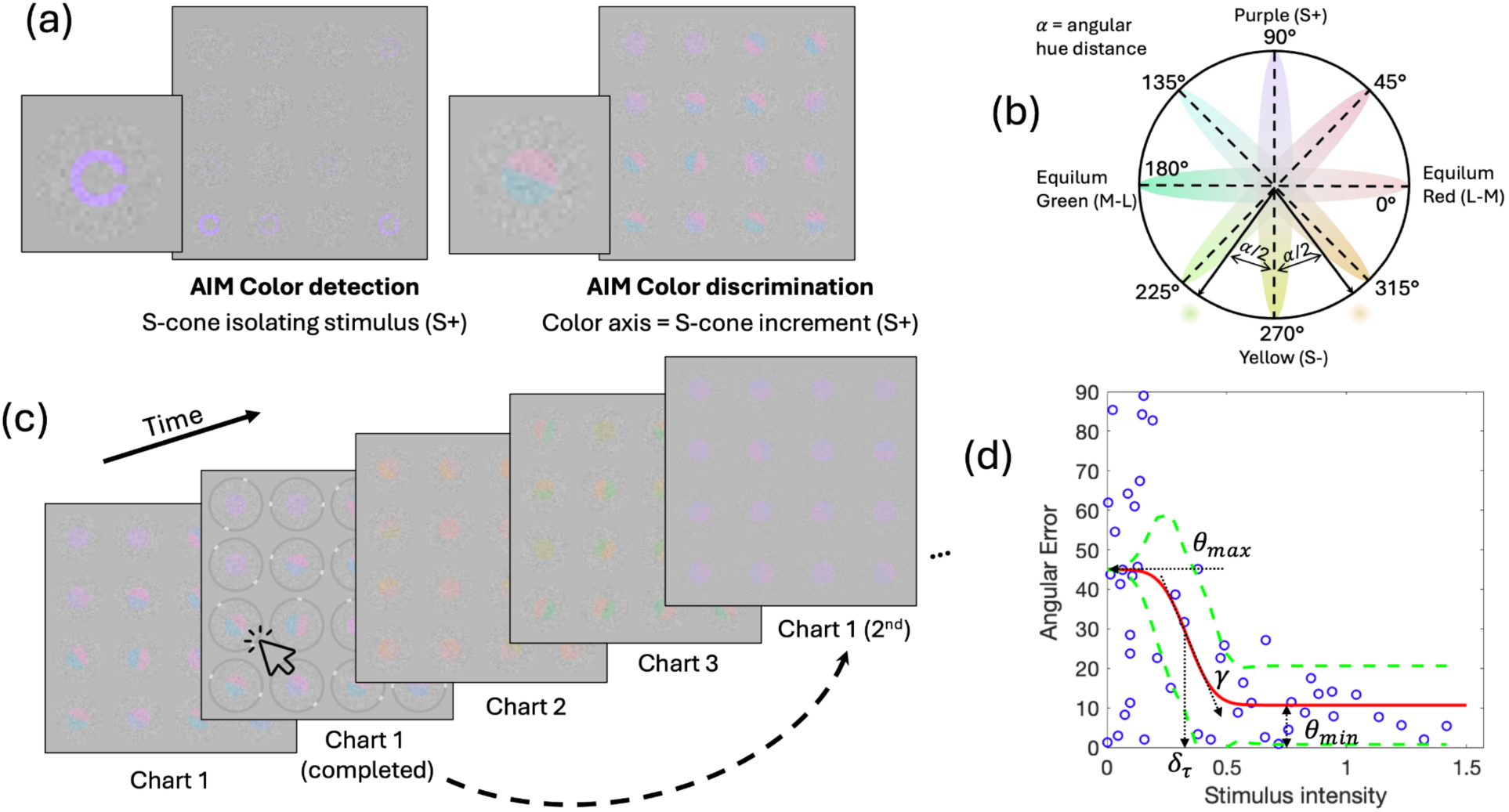
Demonstration of Angular Indication Measurement (AIM) stimuli and experimental procedures. (a) AIM Color Detection (left; S-cone isolating stimuli) and Discrimination (right; stimuli around S-cone increment direction) task interfaces. An enlarged picture of the bottom left cell of each interface is shown. (b) A color wheel showing the equiluminance plane with four primary axes representing stimuli for stimulating the red-green and blue-yellow post-receptoral mechanisms as in the DKL color space (Derrington, Krauskopf, & Lennie, 1984), as well as four intermediate axes. Eight color directions in total were tested in the discrimination task. For example, for the yellow (S−, 270°) discrimination condition, two colors were presented at equal symmetric angular distances (α/2) from yellow. The colors are an illustration but not the actual colors used for generating the stimuli. (c) AIM experimental procedures illustrated for color discrimination. The dashed arrow indicates that the procedure is adaptive: the range of stimulus pairs on the second chart is based on analysis of the responses to stimuli on the first chart. (d) An example of a typical psychometric function: blue circles represent the absolute angular error of each selection plotted against stimulus intensity (cone contrast for detection; color difference angle for discrimination); the red curve depicts the best-fitting function for Equation 1 and the green dashed lines indicate the upper and lower 95% confidence intervals. Estimated parameters are denoted by dotted arrows.

This article has five goals. First, we introduce the AIM Color detection and discrimination method, and explain the features. Second, with the AIM threshold data collected across color normal (CN) and CVD groups, we use supervised machine learning clustering tool to aid the categorization of CVD subtype categories. Third, we compare AIM categorization results against Rayleigh matches on an anomaloscope, the HRR plates (4^th^ Edition), and the Farnsworth–Munsell 100 hue tests (FM100). We have previously introduced a related but fundamentally different paradigm named FInD (He, Bex, et al., 2023). FInD utilizes an adaptive *d-prime* method to estimate detection and discrimination thresholds and requires participants to click on visual stimuli that are visible or appear different to them. We also compare AIM Color Detection and Discrimination tasks to the FInD Color Detection and Discrimination tasks in the Discussion section. Fourth, we examine the test-retest repeatability for the AIM thresholds. Lastly, we report AIM’s additional outcomes.

## Methods

### Participants

Twenty-three self-identified CN participants (mean ± SD age: 26 ± 8; age range: 19-56; 14 females) and fifteen self-identified CVD participants (mean ± SD age: 24 ± 9; age range: 19-52; 2 females) were recruited. All had normal or corrected-to-normal visual acuity as indicated using the AIM Visual Acuity task (Skerswetat et al., 2024), and self-reported to have no history of eye diseases except for one CN participant who had strabismus and amblyopia. Participants with refractive error wore their own corrective lenses throughout the experiment. All participants provided informed consent and filled out a questionnaire inquiring demography and ocular history before the experiment. Approval was granted by Northeastern University Institutional Review Board and the experimental procedures followed the principles of the Declaration of Helsinki.

### Apparatus

MATLAB (MathWorks, Natick, MA) was used to generate experimental procedures (R2021a) and perform data analysis. Stimuli were created by Psychtoolbox (Kleiner, Brainard, & Pelli, 2007), and presented on an HP all-in-one computer, with display resolution 3839×2159 pixels (60.2° × 36.4° at 60 cm) and 60Hz refresh rate. The display was calibrated and gamma corrected with a Photo Research PR-670 spectroradiometer (Photo Research, Chatsworth, CA). Luminance of the mid-grey background was 130.6 cd/m^2^. Participants viewed the screen using both eyes with head position fixed by a chin rest at a distance of 60 cm. Rayleigh matches were completed on an Oculus HMC anomaloscope (Oculus, Germany). Illumination (1069 cd/m^2^) for HRR and FM100 administration was provided by a Sol•Source daylight lamp (117V, 50/60 Hz) manufactured by GretagMacbeth. Test durations for HRR, FM100, and anomaloscope measurements were recorded with a mobile phone timer application or by MATLAB (MathWorks, Natick, MA) for AIM tasks.

### Tasks and stimuli

Rayleigh color match, Hardy-Rand-Rittler (HRR) Pseudoisochromatic Plates (4th Edition), Farnsworth-Munsell 100 hue test (FM100), FInD Color Detection and Discrimination tasks (He, Bex, et al., 2023), and AIM Color Detection and Discrimination tasks were assessed in this study.

#### Rayleigh Color Match

Participants were asked to complete four Rayleigh matches with each eye. Additionally, CVD participants completed one more trial to indicate their matching range. The AQ matching range was established by having participants identify the leftmost (highest anomalous quotient) and rightmost matches (lowest anomalous quotient). The experimenter chose a variety of stimuli covering a wide range of color pairs, and CVD participants were instructed to describe the color appearance of the stimulus and indicate whether the top and bottom colors matched in terms of brightness and hue until they perceived the colors as matching. Anomalous quotient scores for the Raleigh matches, CVD color matching range, and test duration were recorded.

#### HRR Pseudoisochromatic Plates

Participants were asked to report the shape and location of color symbols on each plate while the experimenter flipped through the pages. The number of mistakes on each plate and the test duration were recorded.

#### FM100

Participants were instructed to order the 85 caps according to hue for all testing cases. The four cases were completed in random order. The response scores and the test duration were recorded.

#### FInD Color Detection and Discrimination

The same protocol as reported in He, et al. (2023) was deployed for this study.

#### AIM paradigm

The AIM paradigm presented one chart at a time, each containing 4×4 cells (Figure 1a). A stimulus, either a 2°⌀ Landolt C (detection) or a 2°⌀ bipartite circular patch (discrimination), was presented in each 5° cell and embedded in 14 Hz, 20% contrast dynamic luminance noise with check size 3.8 arcmin. The orientation (the gap of the Landolt C for detection, or the central edge for discrimination) and the signal-intensity of each stimulus (cone contrast for detection, or color difference for discrimination) were randomized. Thus, AIM does not suffer from memory biases of printed, fixed range tests. Each task presented three charts for each color condition (3 cone increment conditions for detection, or 8 testing color directions for discrimination), with later charts responsively adaptive to previous charts (Figure 1c). Specifically, all existing responses were fit with a cumulative Gaussian psychometric function (see Equation 1). The estimated ±95% confidence interval for threshold was used to select the range of stimulus intensities for the next chart.

#### AIM Color detection

AIM Color Detection stimuli were Landolt Cs (2°, Figure 1a, left) with L-, M-, or S-cone isolated colors. Details regarding computation of cone isolating directions can be found in He, Taveras-Cruz, & Eskew (2021). Stimulus contrast was adaptively controlled by the AIM algorithm. Cone contrast detection thresholds for L, M and S colors and test duration were recorded.

#### AIM Color discrimination

AIM Color Discrimination stimuli were 2°⌀ circular patches in which a central blurred (σ=0.1°) edge separated the patch into two semicircular halves. The two halves contained different colors around one of eight testing colors (Figure 1a, right). Testing colors were selected from an equiluminant color plane with the four orthogonal directions being equiluminant red, equiluminant green, S-cone incremental, and S-cone decremental axes, probing the post-receptoral red-green and blue-yellow channels. Four intermediate equiluminant color directions were also selected between the four primary axes (Figure 1b). Color contrast was fixed at 6×detection threshold, so all stimuli are equally detectable. In each stimulus, the two colors in the semicircles were at the same angular distance (α) away from the testing color direction. The threshold angular distance around each testing color direction defined color discriminability. The angular distance between the two colors was adaptively controlled by the AIM algorithm (Figure 1b). Color discrimination thresholds around 8 color directions and test duration were recorded.

### Procedures

Participants’ vision was screened before the experiment, including visual acuity and autorefraction, to ensure best corrected visual acuity ≥ 20/40. Each participant completed the color tasks in random order either once (test session) or twice (test-retest sessions) on different days.

For the AIM tasks, participants were instructed to indicate the orientation of each C’s gap or the central edge by clicking the corresponding location on the surrounding ring. The difference between indicated angle and the true angle is recorded as angular error (θ*_err_*). A psychometric function can then be constructed from the angular error as a function of stimulus intensity (cone contrast for detection, or color difference angle for discrimination; Equation 1; Figure 1d). Three parameters were estimated directly from Equation 1 for fitting this function, including detection or discrimination threshold (δ_τ_), slope (γ), and noise (report error for highly visible stimuli, θ*_min_*):

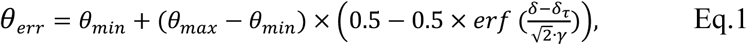

where δ is either cone contrast (detection) or color difference angle (discrimination), and θ*_max_* is the orientation error for a guess response (θ*_max_* = 90° for detection; and 45° for discrimination). The estimated parameters (δ_τ_, γ, and θ*_min_*) are constrained in a reasonable range to determine thresholds. MALTAB functions *fittype*, *fit*, and *predint* are used to fit Equation 1 and the confidence interval to the data.

## Results

The main analyses are based only upon the test session data except for the repeatability analysis. Descriptive statistics of FM100 and AIM tasks for CNs are reported in Supplementary Materials (Table S1).

### Rayleigh Match

The anomalous quotient (AQ) of all CNs scored between 0.7 and 1.4 and were therefore diagnosed as normal according to the criteria supplied with the Oculus HMC anomaloscope. The type and severity of the anomaloscope testing is summarized in Table 1 for each CVD. Details of the CVD AQ range and matching values can be found in Table S2. The average time to complete the task was 14.84 min (median 13.02 min) for CNs to complete eight matches and 13.87 min (median 13.83 min) for CVDs to complete eight matches with an additional matching range trial.

**Table 1.** Comparison of anomaloscope color matching results with SVM classification scores for AIM results. Stage 1 classification scores are the “CVD group” scores and stage 2 classification scores are the “deutan group” scores. Positive number values indicate that they fall in the current group whereas negative values suggest it is classified in the alternative group.

| CVD# | Rayleigh Match |  | Stage 1 classification scores ("is CVD" scores) |  |  |  |  |  | Stage 2 classification scores ("is deutan" scores) |  |  |  |  |  |
| --- | --- | --- | --- | --- | --- | --- | --- | --- | --- | --- | --- | --- | --- | --- |
|  | Type | Severity | 1 | 2 | 3 | 4 | 5 | AVG | 1 | 2 | 3 | 4 | 5 | AVG |
| 1 | Protan | simple | 1.33 | 1.00 | 0.83 | 1.00 | 1.00 | 1.03 | -0.04 | -1.00 | -1.00 | -1.00 | -1.00 | -0.81 |
| 2 | Protan | dichromat | 1.00 | 1.01 | 1.00 | 1.09 | 1.13 | 1.05 | -1.00 | -1.00 | -0.57 | -1.00 | -1.00 | -0.91 |
| 3 | Deutan | simple | 1.43 | 1.13 | 1.25 | 1.17 | 1.10 | 1.21 | 1.05 | 1.00 | 1.14 | 1.10 | 1.10 | 1.08 |
| 4 | Deutan | simple | 1.00 | 1.00 | 1.00 | 1.00 | 0.86 | 0.97 | 1.00 | 0.44 | 1.00 | 1.00 | 1.00 | 0.89 |
| 5 | Deutan | dichromat | 3.22 | 2.82 | 2.94 | 2.81 | 2.72 | 2.90 | 1.19 | 1.16 | 1.27 | 1.26 | 1.24 | 1.23 |
| 6 | Protan | dichromat | 1.69 | 1.61 | 1.49 | 1.54 | 1.77 | 1.62 | -1.00 | -1.23 | -1.10 | -1.43 | -1.46 | -1.24 |
| 7 | Deutan | extreme | 5.85 | 6.78 | 6.51 | 6.21 | 6.58 | 6.39 | 1.62 | 1.46 | 1.00 | 1.00 | 0.72 | 1.16 |
| 8 | Deutan | simple | 0.17 | 0.87 | 0.83 | 0.72 | 0.81 | 0.68 | 1.00 | 1.00 | 1.28 | 1.13 | 1.09 | 1.10 |
| 9 | Protan | extreme | 2.38 | 1.96 | 1.88 | 1.94 | 2.25 | 2.08 | -1.11 | -1.35 | -1.41 | -1.74 | -1.78 | -1.48 |
| 10 | Deutan | extreme | 2.89 | 3.15 | 3.14 | 3.02 | 2.97 | 3.03 | 1.24 | 1.00 | 1.31 | 1.29 | 1.22 | 1.21 |
| 11 | Deutan | extreme | 6.47 | 7.19 | 7.07 | 6.65 | 6.93 | 6.86 | 1.75 | 1.99 | 1.87 | 1.82 | 1.59 | 1.80 |
| 12 | Protan | dichromat | 3.00 | 2.62 | 2.42 | 2.57 | 2.82 | 2.69 | -1.27 | -1.93 | -2.07 | -2.32 | -2.38 | -1.99 |
| 13 | Deutan | extreme | 4.58 | 3.16 | 3.39 | 3.27 | 3.24 | 3.53 | 1.00 | 0.96 | 1.00 | 1.00 | 1.00 | 0.99 |
| 14 | Protan | extreme | 4.05 | 3.79 | 3.41 | 3.61 | 4.04 | 3.78 | -1.59 | -2.42 | -3.17 | -3.39 | -3.54 | -2.82 |
| 15 | Deutan | simple | 1.00 | 1.00 | 1.00 | 0.83 | 1.00 | 0.97 | 1.24 | 1.20 | 1.22 | 1.15 | 1.10 | 1.18 |

### HRR

CNs and CVDs completed the 24 plates in HRR. CN participants reported all plates correctly and all CVDs made mistakes in the diagnostic plates. For CNs, the average testing duration was 46 seconds (median= 47.5 sec; n=22) for the first 10 plates and 1.41 min (median=1.48 min; n=19) for all plates. For CVDs, the average testing duration was 2.78 min (median=2.77 min; n=13) for all plates.

### FM100

We report total error scores (TESs) and right-half midpoint (MP) for the FM100 testing. TES is computed by summing the error scores for each color, with a deduction of two from each error score, following standard procedure (Farnsworth, 1943). The right-half MP includes the median value of error scores for cap sequence 43–84 and was utilized to determine the defect type. The calculation and classification criteria are the same as in He, Bex, et al. (2023).

For CNs, the mean and standard deviation of the TES are reported in Table S1. Based on the TES, 21.7%, 69.6%, and 8.7% of CN participants were classified as superior, average, and low discrimination ability, respectively. The average error score pattern alongside the standard error range is illustrated in Figure 2a. Typically, CNs demonstrate relatively minimal error scores across hues, consistent with previous findings (He, Bex, et al., 2023; Knoblauch et al., 1987), and can be represented on the plot using a maximum error score scale of 4. In contrast, CVDs exhibited higher TES values with 80% classified as having low discrimination ability. The remaining three participants were classified as average ability. An example of a CVD error score pattern is presented in Figure 2b. Notably, compared to the CN error pattern, the CVD error pattern demonstrates a more pronounced and spiky nature with higher magnitudes (note different axis ranges for CN and CVD data). Error patterns for each CVD are shown in Figure S1. In line with prior research (Birch, 1989; He, Bex, & Skerswetat, 2023), the TES values of four CVDs fall within the error score range observed for CNs, showing better discrimination ability than some of the CNs.

**Figure 2.**
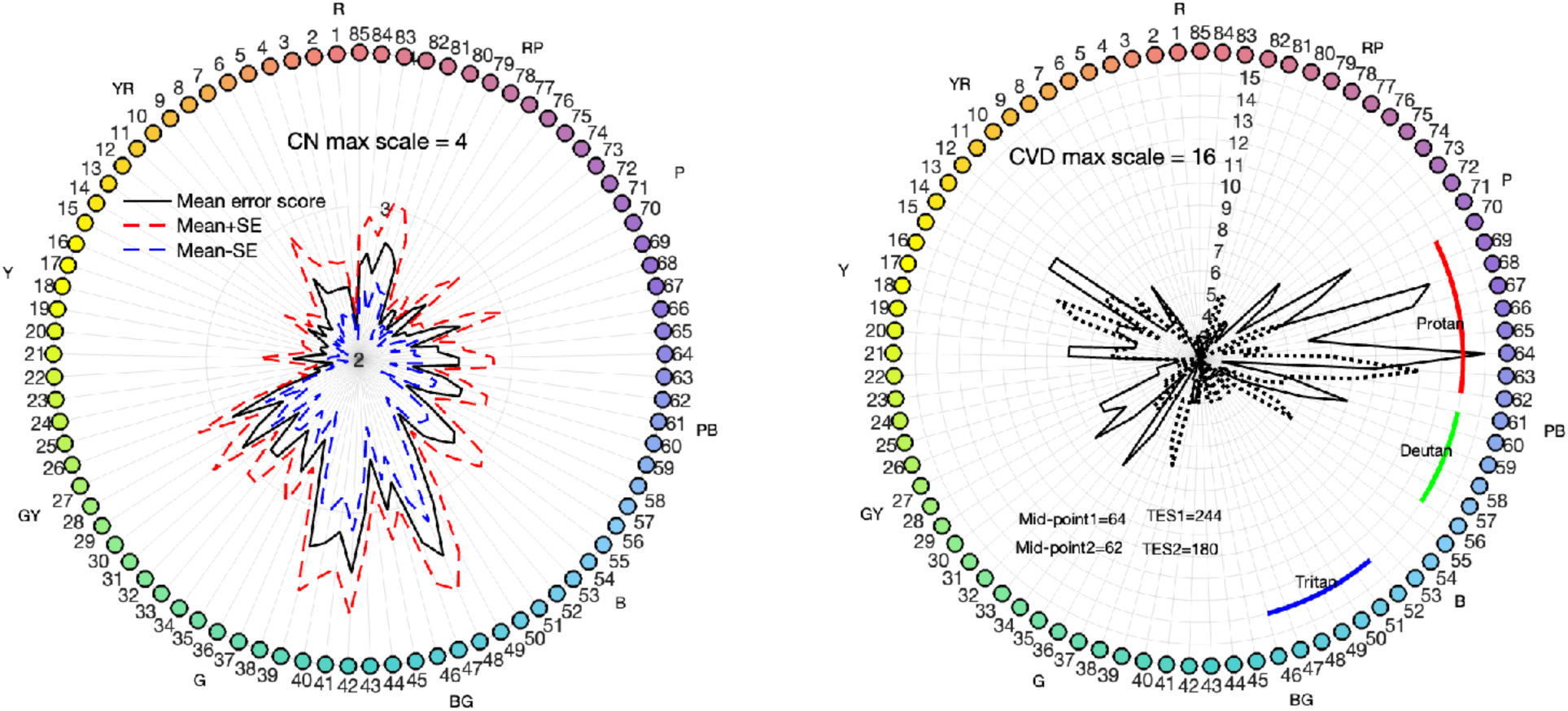
FM100 test results. Left: Average error pattern for CNs. Each hue of the colored caps is numerically designated from 1 to 85. The solid black curve delineates the average error score pattern for 23 CNs. The standard error range is depicted by red and blue dashed lines, indicating the upper and lower bounds, respectively. Note that the range of the radial scale is 2 to 4. Right: the error score pattern from a representative CVD individual (CVD#12). The solid and dotted curves represent test and retest data, respectively. Note that the range of the radial scale is 2 to 15. Additionally, the midpoint and total error score of this participant are indicated in the figure. Note that the midpoint of this individual falls within the protan range for both test and retest sessions.

Classification of the CVD type was conducted according to the right-half MP values. The classification criteria are outlined in the right panel of Figure 2, delineating the MP ranges for protans, deutans, and tritans as 62–70, 56–61, and 46–52, respectively. The TES and MP scores for the color vision deficient (CVD) group are provided in Table S3. The average time was 15.30 min (median 13.60 min) for all CNs to complete eight matches and 13.84 min (median 14.00 min; n=14) for CVDs to complete eight matches with an additional matching range trial.

### AIM Color Detection and Discrimination

AIM Color detection and discrimination thresholds estimated from psychometric functions and rescaled by vector proportions for the detection thresholds are displayed in the top and bottom panels of Figure 3, respectively. The 23 colored circles for each testing direction represent CN reference data while the black crosses show the color thresholds of each CVD participant in each panel. The anomaloscope classification for each CVD is added to each title. For both tasks, thresholds of CNs tend to cluster at lower values, while thresholds of CVD observers are selectively elevated and sometimes outside the range defined by CN thresholds. Protan and deutan participants show distinct elevation patterns. Protans have highly elevated L-cone thresholds compared to CNs and deutans, furthermore their M-cone thresholds are slightly but significantly elevated compared to CNs. Deutans show the opposite pattern, with M-cone thresholds elevated most and L-cone thresholds second most (Figure 5). Both CVD groups show comparable discrimination threshold elevation in Purple and Yellow testing directions (Figure 6).

**Figure 3.**
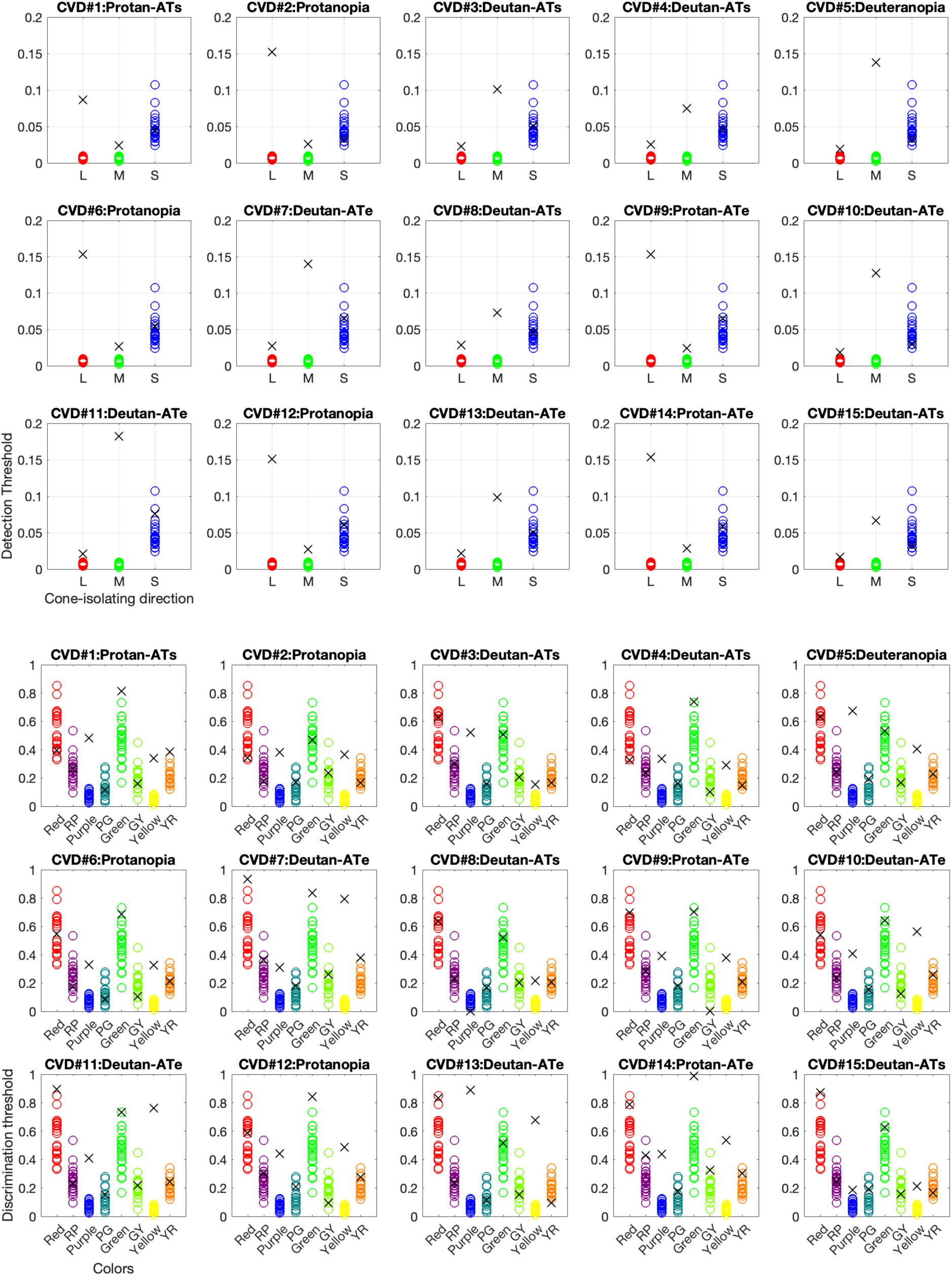
AIM Color detection (top panels) and discrimination (bottom panels) thresholds estimated from the psychometric functions for each CVD participant. The detection thresholds are in arbitrary cone contrast units and the discrimination thresholds are in radian units. Each panel represents one CVD participant’s data (black crosses) compared with the dataset for all 23 CNs (colored circles). Above each panel, the letter following the dash indicates their deficiency severity, where “AT”, “s” and “e” represent “anomalous trichromat”, “simple” and “extreme”, respectively.

Two-way ANOVA examining detection thresholds of CN, protan, and deutan CVD groups, across testing colors, showed significant main effects of group and color, as well as a significant interaction between the two for both protan (group: *F(1,81)=417.02, p<.001*; color: *F(2,81)=80.43, p<.001*; interaction: *F(2,81)=127.18, p<.001*) and deutan (group: *F(1,90)=403.21, p<.001*; color: *F(2,90)=42.81, p<.001*; interaction: *F(2,90)=148.04, p<.001*) groups. For the discrimination task, CN and either protan (group: *F(1,216)=50.5, p<.001*; color: *F(7,216)=34.1, p<.001*; interaction: *F(7,216)=21.57, p<.001*) or deutan (group: *F(1,240)=60.7, p<.001*; color: *F(7,240)=39.31, p<.001*; interaction: *F(7,240)=15.42, p<.001*) groups are significantly different in thresholds for purple and yellow color directions. These results underscore the importance of the L, M-cone detection and the purple and yellow discrimination conditions over other color conditions tested for diagnosing protan and deutan deficiency types.

Additionally, significant differences between CN and the two CVD groups were observed for noise and slope in L and M detection conditions and Purple and Yellow discrimination conditions. Complete analyses for all parameters are reported in the Supplementary material (Tables S4, S5).

On average, CN participants took 6.39 ±2.78 minutes (average: 42.62 seconds/chart) to complete all detection task trials (3 color conditions ×3 charts;), while CVD participants took 4.85 ±1.28 minutes (average: 32.36 seconds/chart). For the discrimination task, CN participants took 16.56 ±7.29 minutes (average: 41.39 seconds/chart), and CVD participants took 13.38 ±3.57 minutes (average: 33.44 seconds/chart; 8 color conditions ×3 charts).

### Classification of CVD type using AIM thresholds in combination with supervised machine learning

To classify subjects to various color vision type and severity categories, we adopted the Support Vector Machine (SVM) algorithm to train machine learning models for this task. SVM is a robust tool for tackling classification problems and is extensively used across various fields including medicine for disease classification (Pisner & Schnyer, 2020). In our case, AIM detection and discrimination thresholds together provide 11 features (each color condition is treated as one feature), with anomaloscope color-matching diagnosis (either CN or CVD and either protan or deutan) serving as the ground-truth labels for each participant. The dataset comprised 38 data points for each feature from both CN and CVD participants. Given such a small dataset, the cross-validation procedure is needed to minimize bias in selection of the training and testing dataset. Five-fold cross-validation was implemented, which involves partitioning the dataset into five folds, and using each fold in turn as the test set while the remaining four folds serve as the training set.

We performed a two-stage SVM classification. The first stage trained a binary SVM classifier to differentiate CNs and CVDs, with the classification scores retained for further severity estimation. These scores, which ranged from-∞ to ∞, indicate the distance of a data point from the decision boundary set by the classifier. In binary classification, the classification scores reflect the degree of affiliation of the AIM thresholds with either group: positive values suggest membership in the target group, while negative values indicate the opposite. We utilized these distance scores to estimate how different the data points of CVDs are from the CN groups and thus estimate the severity of color deficiency. The second stage trains another binary SVM classifier to distinguish between protan and deutan deficiency types (see details in Supplementary Materials).

We found that the first stage of SVM classification using a polynomial kernel achieved an average cross-validated accuracy of 100%, effectively distinguishing between CNs and CVDs. The second stage repeated the procedure but focused only on the threshold data from 15 CVDs and still achieved 100% accuracy. These perfect accuracies suggest robust performance of SVM classifiers for identifying whether the thresholds reflect normal color vision, and if not, whether it is a protan or deutan deficiency. The classification scores are continuous in nature and we subdivided these scores into “simple”, “extreme” and “dichromat” groups as defined by the anomaloscope outcomes. The 5-fold cross-validation yielded five trained models and corresponding sets of scores. These scores are averaged and reported in Table 1. All AIM data classified as “simple” have lower scores ranging from 0.68 to 1.21, and all “extreme” data have higher scores than those of “simple”, ranging from 2.08 to 6.86. However, the “dichromat” scores overlap with the ranges of the “simple” and “extreme” groups, while they are expected to be the highest. One possibility is that the anomaloscope-based ground-truth is inaccurate—extreme anomalous trichromats (AT) can have close matching ranges as dichromats (Birch, 2003). For simplicity, scores less than or equal to 1.21 were classified as anomalous trichromats, and those larger than or equal to 1.62 as dichromats or extreme AT. The final classification results are detailed in Table 2.

**Table 2.** A summary of testing results of each CVD participant for all tasks. The deficiency types are either protan or deutan. The severity categories differ across tasks—Rayleigh match: simple, extreme, or dichromat; HRR: mild, medium, or strong; FM100: superior, average, or low discrimination ability; AIM: simple or dichromat/extreme AT (D/E). The classification that is comparable and different from the anomaloscope classification is underlined and italicized.

| CVD# | Rayleigh Match |  | HRR |  | FM100 |  | AIM-SVM |  |
| --- | --- | --- | --- | --- | --- | --- | --- | --- |
|  | Type | Severity | Type | Severity | Type | Severity | Type | Severity |
| 1 | Protan | simple | <u>Deutan</u> | Mild | Protan | Low | Protan | simple |
| 2 | Protan | dichromat | Protan | Medium | Protan | Low | Protan | <u>simple</u> |
| 3 | Deutan | simple | Deutan | Medium | <u>Protan</u> | Low | Deutan | simple |
| 4 | Deutan | simple | <u>Protan</u> | Strong | <u>Protan</u> | Average | Deutan | simple |
| 5 | Deutan | dichromat | Deutan | Strong | Deutan | Low | Deutan | D/E |
| 6 | Protan | dichromat | Protan | Strong | Protan | Low | Protan | D/E |
| 7 | Deutan | extreme | Deutan | Strong | Deutan | Low | Deutan | D/E |
| 8 | Deutan | simple | Deutan | Mild | <u>Protan</u> | Average | Deutan | simple |
| 9 | Protan | extreme | Protan | Medium | Protan | Low | Protan | D/E |
| 10 | Deutan | extreme | Deutan | Strong | <u>Protan</u> | Low | Deutan | D/E |
| 11 | Deutan | extreme | Deutan | Strong | Deutan | Low | Deutan | D/E |
| 12 | Protan | dichromat | Protan | Strong | Protan | Low | Protan | D/E |
| 13 | Deutan | extreme | Deutan | Strong | Deutan | Low | Deutan | D/E |
| 14 | Protan | extreme | Protan | Strong | Protan | Low | Protan | D/E |
| 15 | Deutan | simple | <u>Protan</u> | Medium | <u>Protan</u> | Average | Deutan | simple |

We also repeated this 2-stage process with reduced features to explore the impact of feature reduction, using four cardinal directions instead of eight color directions for the AIM discrimination task. This adjustment resulted in seven features and maintained 100% accuracy in stage 1 classification. However, in stage 2, the highest accuracy achieved was 93.33% for classifying deficiency type. The lower accuracy in stage 2 can be attributed to the limited sample size and the imbalance among the groups, which complicates achieving perfect and stable performance. Additionally, the thresholds for intermediate color directions, which were excluded in this reduced feature set, may hold valuable information for distinguishing deficiency types.

### Comparison of the methods

Table 2 compares the results of all tests. HRR diagnosis is not very consistent with the anomaloscope diagnosis with three out of fifteen cases showing discrepancies in protan or deutan classifications. The severity types are not directly comparable, as the HRR system categorizes severity only as mild, medium, or strong. It has been reported that the HRR is effective at identifying dichromats but less so for anomalous trichromats (Birch, 1997a). Similarly, FM100 results show five out of fifteen cases that differ from the anomaloscope diagnosis, and the severity types are also not matched to that of the anomaloscope. Poor differentiation between dichromats and ATs has been reported (Birch, 1989; Lakowski, 1969). These results agree with the findings in He, Bex, et al. (2023). Conversely, the AIM Color detection and discrimination thresholds combined with the SVM classification replicates the anomaloscope’s protan/deutan classification and provide a severity classification that agrees 93.3% of the cases, with only one case assigned to a different severity category. The high agreement between AIM and the anomaloscope results can be partially explained by the use of Rayleigh color matching outcomes as training labels. The perfect accuracy reveals that our threshold measurements veridically represent the data structure of the anomaloscope color matching outcomes.

### Test-retest repeatability

Twelve CN and thirteen CVD participants (6 protans and 7 deutans) completed the retest session. All CNs passed HRR again. Bland-Altman analyses (Bland & Altman, 1999) were performed to assess test-retest repeatability. MATLAB (MathWorks, Natick, MA) function *normplot* was used to check whether the thresholds data are normally distributed. Detection thresholds are log transformed to achieve normality. In the Bland-Altman plots in Figure 4, the mean test-retest difference (bias, or <u>d</u>) for all scores and thresholds are close to zero (differences between dotted and solid black horizontal lines) and within the 95% confidence interval (CI) range of bias, indicating negligible test-retest differences. Almost all data points are within the 95% CI range of the limits of agreement (LoA) range. The upper and lower 95% LoA are computed as <u>d</u>±1.96×standard deviation of the mean difference. The data points do not vary with the test-retest mean and the slopes of the best linear fits are not significantly different from 0 (Table 3). All data points cluster together, with the CVD data (yellow circles) spread out more than the CN data (green squares). In sum, the Bland-Altman analysis shows good reliability for AIM Color Detection and Discrimination thresholds.

**Figure 4:**
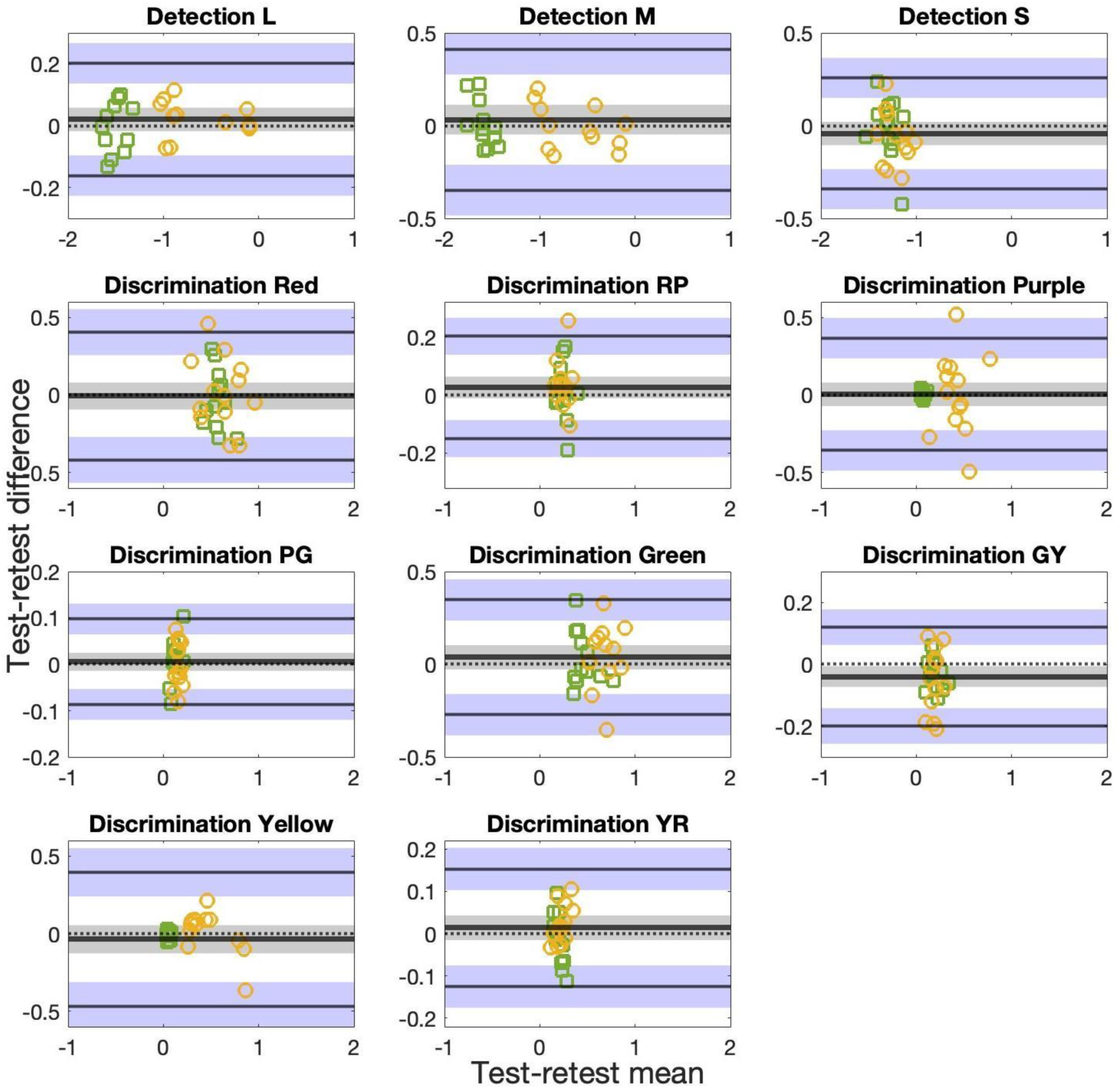
Bland-Altman plots for AIM Color detection L-, M-, and S-cone isolating thresholds and AIM Color discrimination Red (R), Red-Purple (RP), Purple (P), Purple-Green (PG), Green (G), Green-Yellow (GY), Yellow (Y), Yellow-Red (YR) thresholds in radian for 12 CN (green squares) and 13 CVD (orange circles) participants. The thicker solid black line in the middle indicates mean of the test-retest differences (<u>d</u>) and the middle dotted black line indicates a difference of 0 as a reference. These two lines overlap in some panels. Limits of agreement (LoA) lines are depicted as upper and lower black lines. Grey and blue zones are 95%-CI ranges of <u>d</u> and LoAs, respectively, indicating precision of the estimates. Note that scales are selected for each panel to best show data points.

**Table 3:**
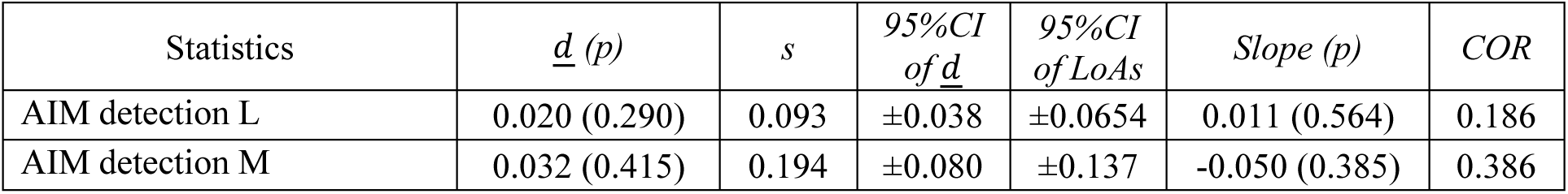
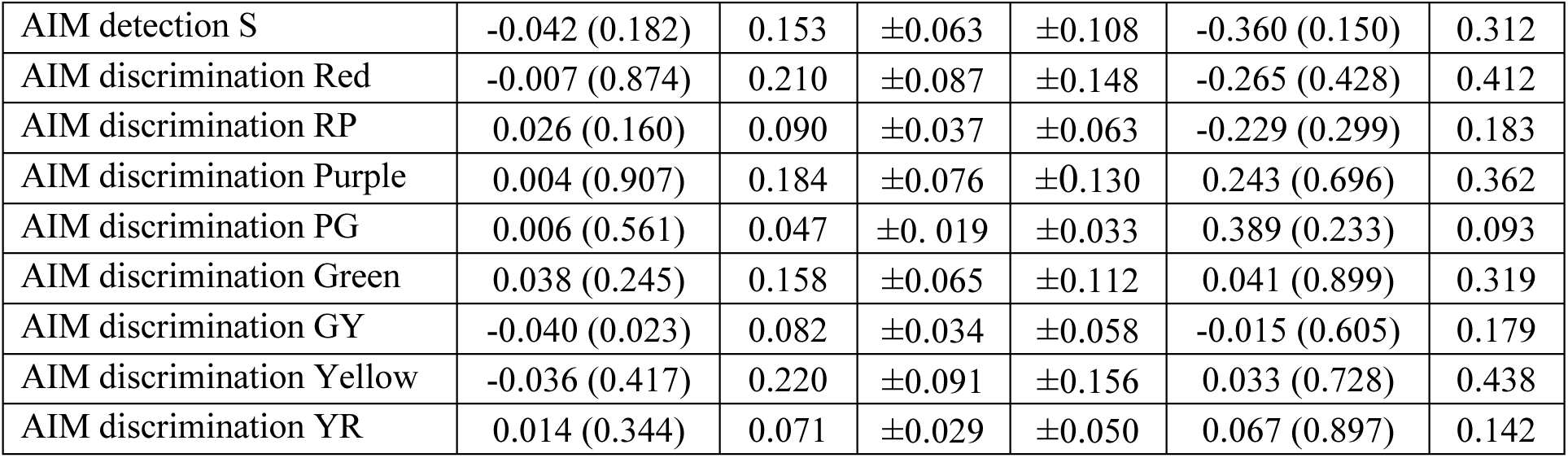
Statistics of the Bland-Altman analysis results for 25 participants. Each column, from left to right, is task, mean test-retest difference or bias (<u>d</u>) and its p value, standard deviation of the differences (s), 95% confidence interval (CI) of <u>d</u>, 95% CI of limits of agreement (LoAs), slope of the best fit line and its p value, and coefficient of repeatability (COR).

## Discussion

We have introduced a novel paradigm, Angular Indication Measurement (AIM), for assessing color vision that enables rapid, self-administered, and personalized interrogation through a response-adaptive approach, while also providing additional diagnostic metrics. The detection and discrimination thresholds derived are sufficiently informative to allow for accurate classification of color vision deficiency types and severity.

### Additional AIM Color parameters

In addition to threshold, we estimated minimum angular error (noise θ_’#$_) and slope (γ) to provide more information from the limited set of data. The recently published article applying the AIM paradigm to visual acuity measurement (Skerswetat, He, et al., 2024) has reported that noise significantly increased with blur and discussed that the AIM acuity psychometric function slope could reflect the progression of eye disease. Tyler (1997) has stressed that psychometric function slopes may affect sensitivity substantially and the steepness is subject to channel uncertainty, spatial and temporal characteristics of the stimuli, and wavelength (Maloney, 1990). In the current study, we have observed significant noise and slope differences across subject groups for selective color directions (Table S5). These results bear more information than a single threshold estimate and hold value for future research.

### Trial-by-trial analysis for AIM charts

Three charts/trials are tested in this study for complete assessment of the AIM tasks, but is three charts per color condition necessary? In Figures 5 and 6 we plot estimated thresholds for using one chart, two charts, and all three charts. Corresponding values are reported in Tables S6 and S7. One way ANOVA shows no significant differences across chart number for each color condition for CN, protan and deutan groups, except for the L-cone detection condition where CNs have chart-one thresholds significantly different from two or three charts (*F(2,66)=5.12, p<.01*). This suggests that two charts, or even one chart, would be sufficient to produce accurate threshold estimation, therefore, less testing time is needed.

**Figure 5.**
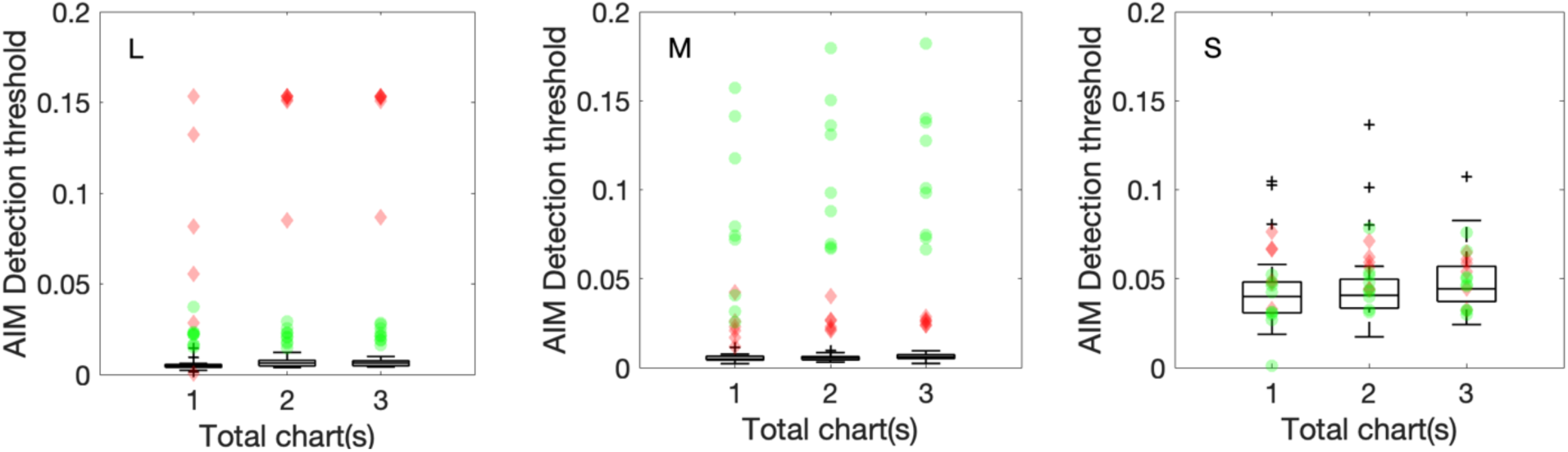
AIM Color Detection thresholds plotted for one chart, two charts, and three charts. Color conditions are L, M, and S-cone isolation from left to right. CN data are represented by the box-whisker plot while protan and deutan data are represented by red diamonds and green circles respectively. In each boxplot, the central line indicates the median, while the lower and upper boundaries show the 25th and 75th percentiles, respectively. Whiskers stretch to the furthest data points that are not outliers, and outliers, values that are more than 1.5 times the interquartile range from the nearest boundary of the box, are marked with black crosses.

**Figure 6.**
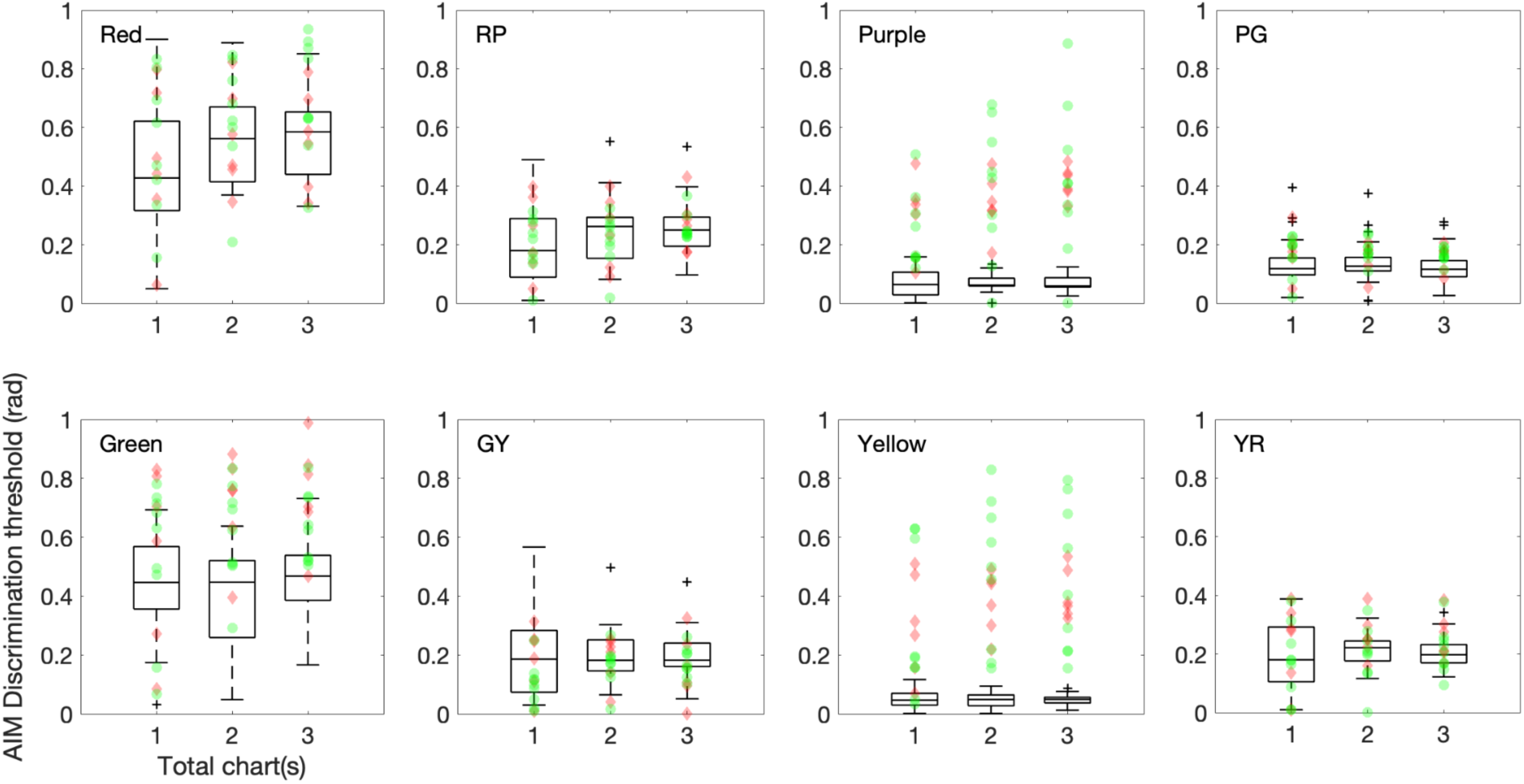
AIM Color Discrimination thresholds plotted for one chart, two charts, and three charts. Color condition is noted on the top left corner in each panel. Symbols are the same as in Figure 5.

### AIM and FInD comparison

The AIM threshold patterns for CNs and CVDs are expected and agree with those reported in He, Bex, et al. (2023). For the detection task, the “S” thresholds for CVDs remain within the CN range, whereas the “L” and “M” thresholds are notably elevated with protans or deutans having significantly elevated “L” or “M” thresholds, respectively, with the other condition slightly elevated as well. For the discrimination task, the thresholds for “Purple” and “Yellow” are significantly increased for CVDs, with occasional increases in “Green” and “Red” thresholds as well.

In He, Bex, et al. (2023) we introduced the FInD paradigm for color assessment. The FInD paradigm utilizes *d-prime* as the stimulus selection criterion and asks the participants to choose the stimuli that appear visible to them; in the current study the AIM paradigm was instead employed as a forced-choice method as participants had to report the orientation in each cell, guessing if necessary, before finishing a chart. A cumulative Gaussian function was then used to estimate threshold and other parameters. Forced-choice methods are known to push the performance to the limit thus yielding the most rigorous and veridical threshold estimation (Taveras-Cruz, He, & Eskew Jr, 2022). In the current study, FInD color detection and discrimination thresholds were also collected under identical experimental conditions from the same participant group. The participants were asked to click on the color blobs that appear visible in the FInD Color detection task and to click on the blob pair that appear to have different colors in the FInD Color discrimination task. Three charts and the same color directions were measured. Comparing AIM thresholds (Figure 3) and FInD thresholds (Figures S2, S3), we observe that the AIM detection thresholds are generally slightly higher, whereas its discrimination thresholds are slightly lower than those of FInD. These differences might be due to AIM and FInD stimuli having different characteristics and response methods (participants can see AIM detection stimuli colors but not able to indicate the orientation correctly at certain contrasts and can compare AIM discrimination stimuli side-by-side but this is not the case for FInD), but the overall threshold patterns of the two paradigms remain consistent.

In research and clinical practices, the AIM and FInD paradigms each have their own strengths. FInD uses a pure signal detection approach and a seen/not seen criterion. In contrast, AIM uses an orientation judgment approach with randomized gap or edge orientations, where the task is to indicate the orientation of a gap of a Landolt C or a color difference of the bipartite stimuli, regardless of the contrast or color difference is above or below threshold. Therefore, AIM and FInD paradigms have different decision criteria. Moreover, as a consequence of its approach, AIM’s psychometric function is a personalized representation of one’s performance. These resulting functions and the additional parameters, as previously discussed, hold promise to be new biomarkers for visual deficits and neuro-ophthalmic diseases.

### SVM classification algorithm

The SVM algorithm’s capability to accurately delineate clear decision boundaries within our dataset suggests that the derived threshold patterns inherently reflect those in the Rayleigh color matches. It is known that severity of inherited color deficiency is continuous instead of discrete (Birch, 1993). With this considered, we classified severity using the distance scores from the stage 1 SVM classifier which is also continuous, and the results showed high concordance with the anomaloscope classifications. This underscores the potential for leveraging the classification score information in severity estimation. However, given the small sample size, the performance of the SVM classifier on a larger dataset or with a different test dataset is not known. Therefore, a model incorporating all collected data and a different testing dataset are required in future research for confirming the high accuracies.

In conclusion, AIM Color detection and discrimination tasks are introduced and validated against HRR, FM100, and anomaloscope color matching diagnosis in this proof-of-concept study. The results exhibit good consistency between the color matching diagnoses and the AIM combined with SVM classifiers, while demonstrating superior testing accuracy of AIM compared to HRR and FM100. Bland-Altman results showed good repeatability and no systematic bias for the AIM color tasks. On average, each AIM chart took around 32-43 seconds to complete. The per-chart analysis suggests that even a single AIM chart per color direction can be sufficient to estimate thresholds, thus reducing the screening time to 2-3 minutes for each AIM task. Small differences in AIM color thresholds relative to our previously introduced FInD paradigm likely arise from variations in experiment protocol, stimulus characteristics, or threshold estimation methods. Overall, AIM proves to be an effective tool for color vision deficiency screening.

## Acknowledgments

This study was supported by NIH grants EY029713 and EY032162. We thank the undergraduate research assistants, Sophia He, Panharath Sok, Launna Atkinson, and Jay Bijesh Shah for helping with data collection and data organization.

