## Supplementary Materials for "Novel color vision assessment tool: AIM Color Detection and Discrimination"

### *Testing results: Tables and figures*

| CN group | FM100 TES | AIM Detection | | | AIM Discrimination (deg) | | | | | | | |
|---|---|---|---|---|---|---|---|---|---|---|---|---|
| | | S | M | L | 0° (R) | 45° (RP) | 90° (P) | 135° (PG) | 180° (G) | 225° (GY) | 270° (Y) | 315° (YR) |
| Test (n=23) | | | | | | | | | | | | |
| Mean (STD) | 49.57 (41.79) | 0.0485 (0.0185) | 0.0061 (0.0016) | 0.0065 (0.0018) | 0.5517 (0.1414) | 0.2568 (0.0924) | 0.0700 (0.0247) | 0.1258 (0.0664) | 0.4622 (0.1324) | 0.1957 (0.0799) | 0.0484 (0.0168) | 0.2088 (0.0519) |
| Retest (n=12) | | | | | | | | | | | | |
| Mean (STD) | 21.33 (18.94) | 0.0445 (0.0107) | 0.0060 (0.0011) | 0.0090 (0.0017) | 0.5491 (0.1175) | 0.2549 (0.0714) | 0.0711 (0.0223) | 0.1322 (0.0624) | 0.4771 (0.1263) | 0.1815 (0.0656) | 0.0468 (0.0192) | 0.2097 (0.0400) |

Table S1. Descriptive statistics of test and retest sessions for the color normal (CN) group.

| CVD# | Type | Severity | AQ range | | | |
|---|---|---|---|---|---|---|
| | | | Min. AQ | Matching | Max. AQ | Matching |
| 1 | Protanomaly | simple | 0.23 | 61.5 (5.5) | 0.26 | 60.1 (6.5) |
| 2 | Protanopia | dichromat | 0.13 | 66.1 (4.4) | Infinity | 0 (32.6) |
| 3 | Deuteranomaly | simple | 2.6 | 23.2 (13.6) | 3.33 | 19.5 (14.3) |
| 4 | Deuteranomaly | simple | 3.08 | 20.6 (14.3) | 3.4 | 19.2 (13.9) |
| 5 | Deuteranopia | dichromat | 0 | 72.7 (12.9) | Infinity | 0 (14.3) |
| 6 | Protanopia | dichromat | 0.02 | 71.8 (2.4) | Infinity | 0 (41.2) |
| 7 | Deuteranomaly | extreme | 0.11 | 66.7 (12.5) | 7.46 | 10.2(13.7) |

| | | | | | | |
|---|---|---|---|---|---|---|
| 8 | Deuteranomaly | simple | 2.2 | 25.9 (10.2) | 3.6 | 18.4 (15.7) |
| 9 | Protanomaly | extreme | 0.34 | 56.9 (11.8) | 24.07 | 3.5 (40) |
| 10 | Deuteranomaly | extreme | 0.52 | 51 (14.1) | 4.73 | 14.9 (15.3) |
| 11 | Deuteranomaly | extreme | 0.16 | 64.7 (13.3) | 8.2 | 9.4 (14.1) |
| 12 | Protanopia | dichromat | 0.09 | 68.2 (3.5) | Infinity | 0 (31.8) |
| 13 | Deuteranomaly | extreme | 0 | 72.9 (13.3) | 4.57 | 15.3 (15.7) |
| 14 | Protanomaly | extreme | 0.14 | 65.5 (4.3) | 31.56 | 2.7 (31) |
| 15 | Deuteranomaly | simple | 4.15 | 16.5 (14.5) | 5.65 | 12.9 (14.5) |

*Table S2. Anomalous quotient (AQ) ranges and their corresponding matching ranges for Rayleigh matches for all CVD participants. Numbers in parentheses represent the corresponding reference light. According to the anomaloscope user's manual (Oculus, Germany), AQ ranging from 0.7 to 1.4 indicate normal matching zone; AQ ranging from less than 0.7 to 0.1 indicate Protanomaly; AQ ranging from larger than 1.4 (mostly larger than 2.0) to infinity indicate Deuteranomaly; AQ up to infinity or down to 0 or including the normal mid-match indicate extreme anomaly. The full range of the mixtures is acceptable for a dichromat. Severity categories are simple, extreme, or dichromat.*

| CVD# | Total error score (TES) | | Mid-point (MP) | |
|---|---|---|---|---|
| | Test | Retest | Test | Retest |
| 1 | 100 | 92 | 71 | 69 |
| 2 | 180 | 152 | 67 | 64 |
| 3 | 168 | 112 | 64 | 66 |
| 4 | 84 | 48 | 70 | 73 |
| 5 | 320 | 252 | 54 | 58 |
| 6 | 216 | 280 | 64 | 59 |
| 7 | 244 | 248 | 60 | 57 |
| 8 | 56 | 72 | 71 | 71 |
| 9 | 316 | 276 | 65 | 59 |
| 10 | 192 | 140 | 63 | 62 |
| 11 | 204 | 172 | 59 | 62 |

| | | | | |
|---|---|---|---|---|
| 12 | 244 | 180 | 64 | 62 |
| 13 | 304 | 296 | 59 | 58 |
| 14 | 176 | 180 | 66 | 64 |
| 15 | 48 | - | 75 | - |

*Table S3. Farnsworth-Munsell 100 hue test (FM100) for color vision deficiency (CVD) observers.*

| Detection | CN (n=15) | | Protan (n=6) | | Deutan (n=9) | |
|---|---|---|---|---|---|---|
| | *F* | *p* | *F* | *p* | *F* | *p* |
| Threshold $\delta_\tau$ | 31.91 | *** | 241.86 | *** | 126.61 | *** |
| Noise $\theta_{min}$ | 5.54 | ** | 4.38 | * | 2.72 | 0.0859 |
| Slope $\gamma$ | 8.51 | *** | 1.88 | 0.1871 | 0.29 | 0.749 |
| Discrimination | CN (n=15) | | Protan (n=6) | | Deutan (n=9) | |
| | *F* | *p* | *F* | *p* | *F* | *p* |
| Threshold $\delta_\tau$ | 114.03 | *** | 6.16 | *** | 5.36 | *** |
| Noise $\theta_{min}$ | 6.77 | *** | 0.96 | 0.4703 | 8.91 | *** |
| Slope $\gamma$ | 7.39 | *** | 0.37 | 0.9127 | 1.21 | 0.3106 |

*Table S4. One-way ANOVA results for AIM Color detection and discrimination threshold, noise, and slope comparing across color directions. All raw data have been log-transformed to achieve normality. P-values <0.001, <0.01, and <0.05 are represented as ***, **, and *, respectively.*

| Detection | Protan | Deutan |
|---|---|---|
| Threshold $\delta_\tau$ | L***, M*** | L***, M*** |
| Noise $\theta_{min}$ | L*** | L**, M*** |
| Slope $\gamma$ | L**, M** | M** |
| Discrimination | Protan | Deutan |
| Threshold $\delta_\tau$ | Purple***, Yellow*** | Purple***, Yellow*** |

| | | |
|---|---|---|
| Noise $\theta_{min}$ | / | Purple***, Yellow*** |
| Slope $\gamma$ | Yellow*** | Purple** |

*Table S5. Two-way ANOVA results for AIM Color detection and discrimination threshold, noise, and slope. This table reports only color conditions where the parameters are significantly different between CN and each CVD group. All raw data have been log-transformed to achieve normality. P-values <0.001, <0.01, and <0.05 are represented as ***, **, and *, respectively. See Tables S6-9 for mean and standard deviation for each group.*

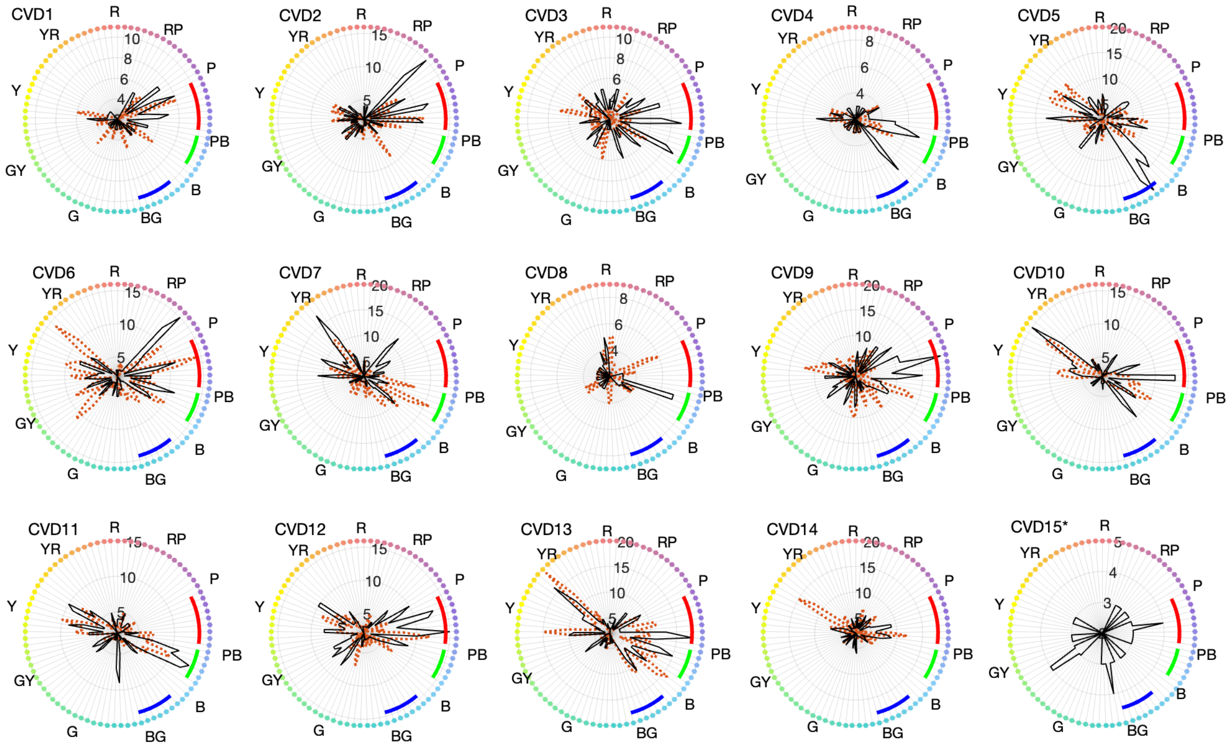

*Figure S1. FM100 error score pattern for 15 CVD observers. Solid black lines and dotted orange lines depict test and retest scores, respectively, for each color cap. The corresponding hues are denoted as color dots around the error score patterns, labeled as follows: R (red), YR (yellow-red), Y (yellow), GY (green-yellow), G (green), BG (blue-green), B (blue), PB (purple-blue), P (purple), and RP (red-purple). The radial scales vary across participants for clarity in visualization of the pattern. Diagnostic criteria for protan, deutan, and tritan are shown as red, green, and blue curve segments, respectively. Participants who did not participate in the retest session is marked with * after the identification number (CVD#15 only).*

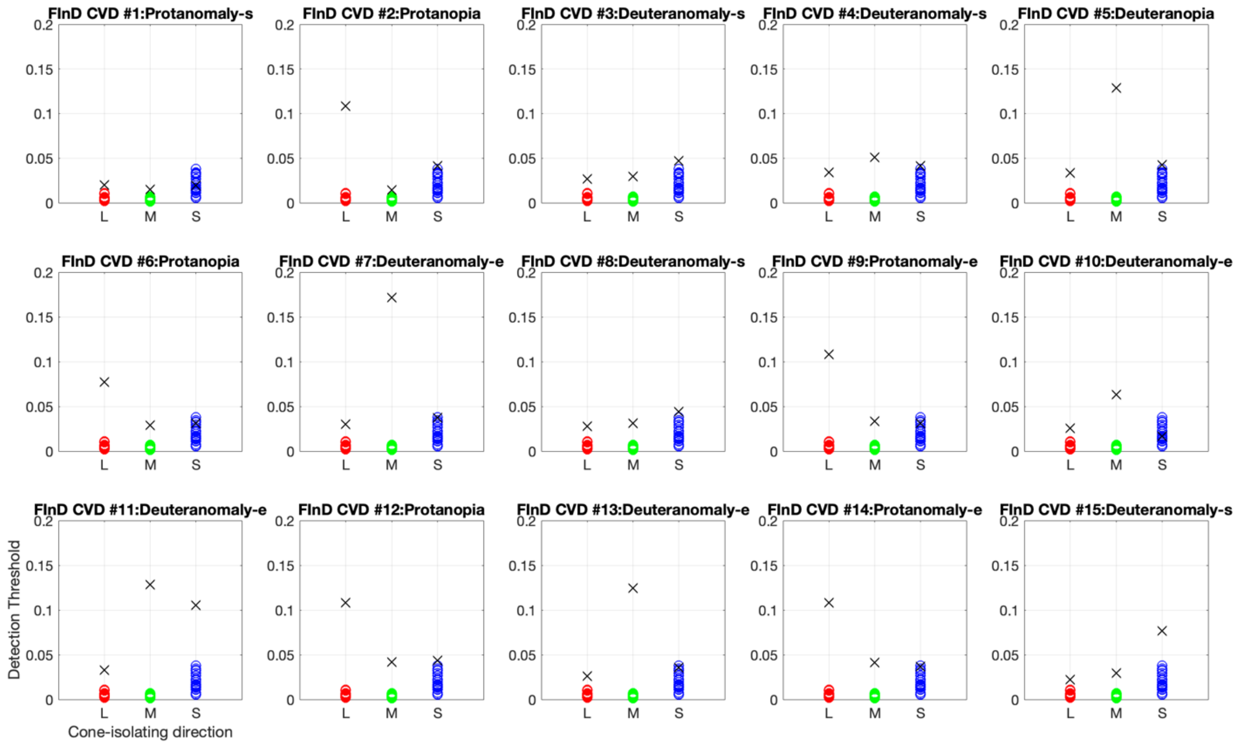

*Figure S2. FInD Color detection thresholds in arbitrary unit. Each panel represents one CVD participant's data (black crosses) compared with the normal dataset (colored circles). Above each panel, the letter following the dash indicates their deficiency severity, where "s" and "e" represent "simple" and "extreme", respectively. Some data points might seem to be at 0 threshold but they are actually all above 0.*

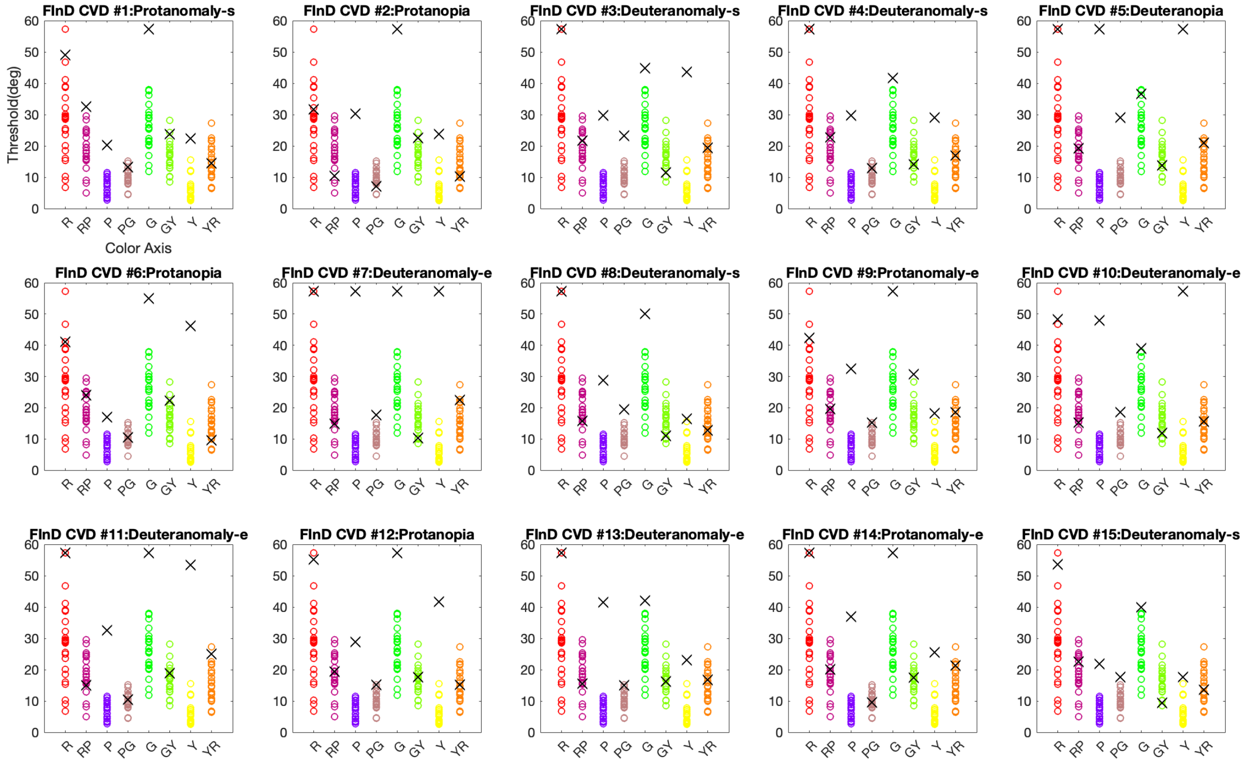

*Figure S3. FInD Color discrimination thresholds in degree unit. Each panel represents one CVD participant's data (black crosses) compared with the normal dataset (colored circles). Above each panel, the letter following the dash indicates their deficiency severity, where "s" and "e" represent "simple" and "extreme", respectively.*

### ***Supervised Machine Learning for classification: Support Vector Machine (SVM)***

MATLAB Statistics and Machine Learning Toolbox was used to perform SVM classification. The cross-validation is achieved by using *cvpartition* class to partition the dataset to 5 folds. For each cross-validated training and testing dataset, function *fitcsvm* was used for training a SVM model, with the kernel being *linear*, *polynomial*, or *gaussian*. Then the trained model is used to predict (function *predict*) the labels and the classification scores.

### ***Trial by trial analysis***

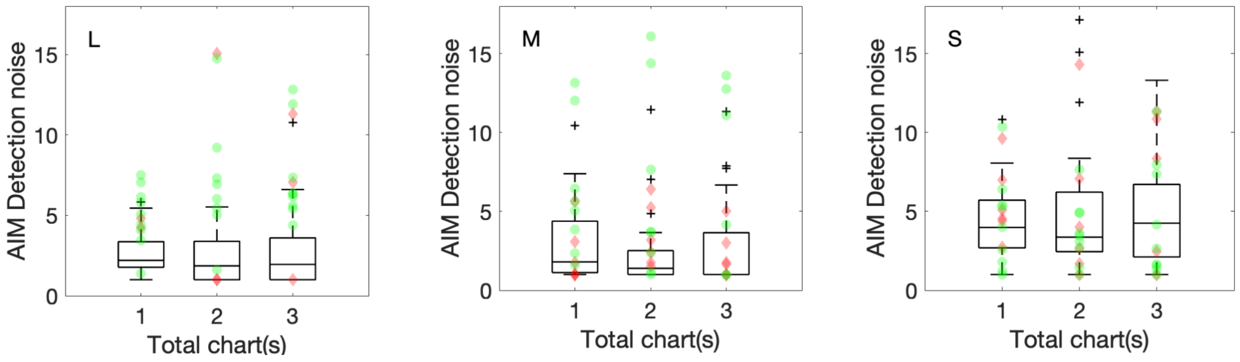

*Figure S4. AIM Color Detection noise plotted for one chart, two charts, and three charts. Color conditions are L, M, and S-cone isolation from left to right. CN data are represented by the box-whisker plot while protan and deutan data are represented by red diamonds and green circles respectively. In each boxplot, the central line indicates the median, while the lower and upper boundaries show the 25th and 75th percentiles, respectively. Whiskers stretch to the furthest data points that are not outliers, and outliers, values that are more than 1.5 times the interquartile range from the nearest boundary of the box, are marked with black crosses.*

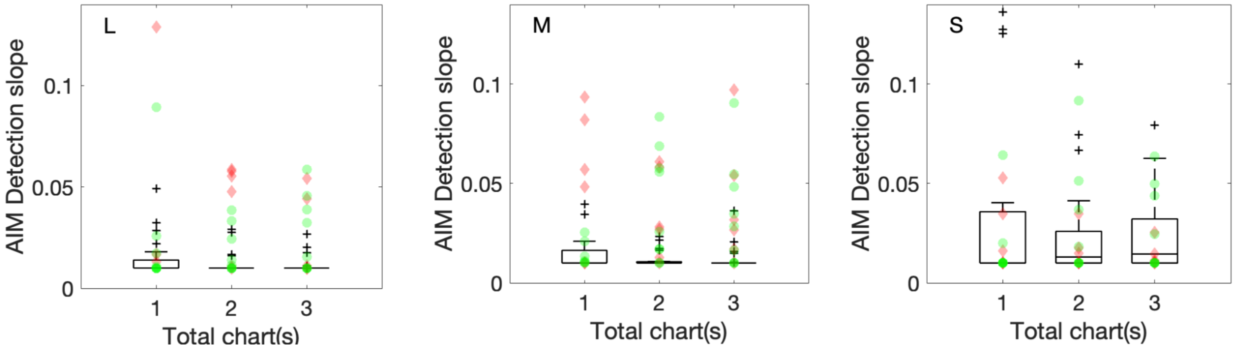

*Figure S5. AIM Color Detection slope plotted for one chart, two charts, and three charts. Symbols are the same as in Figure S4.*

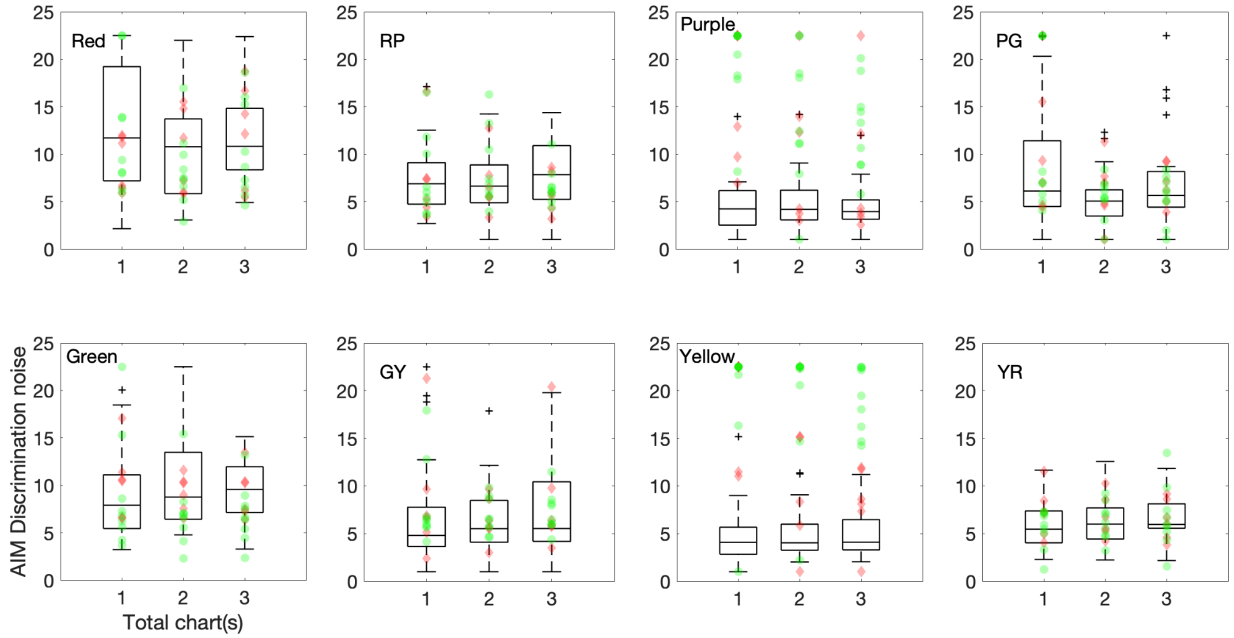

*Figure S6. AIM Color Discrimination noise plotted for one chart, two charts, and three charts. Color condition is noted on the top left corner in each panel. Symbols are the same as in Figure S4.*

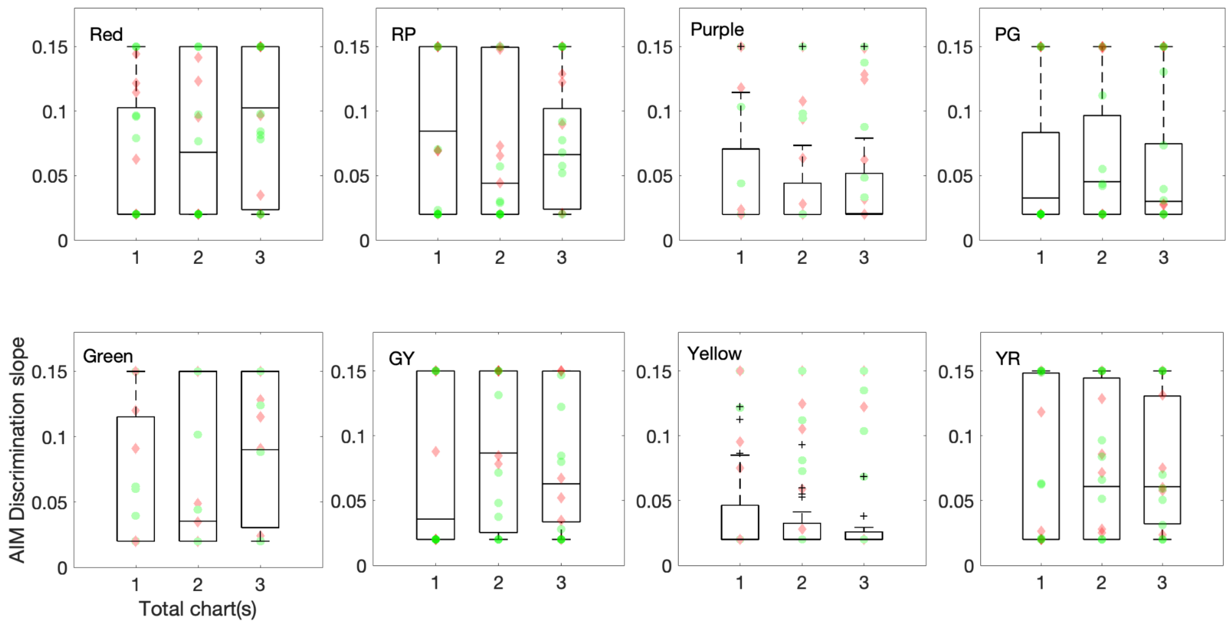

*Figure S7. AIM Color Discrimination slope plotted for one chart, two charts, and three charts. Color condition is noted on the top left corner in each panel. Symbols are the same as in Figure S4.*

| **Threshold** | L | | | M | | | S | | |
|---|---|---|---|---|---|---|---|---|---|
| #of Charts | 1 | 2 | 3 | 1 | 2 | 3 | 1 | 2 | 3 |
| CN | 0.0052 (0.0028) | 0.0065 (0.0019) | 0.0065 (0.0018) | 0.0053 (0.0020) | 0.0057 (0.0016) | 0.0061 (0.0016) | 0.0440 (0.0231) | 0.0477 (0.0263) | 0.0485 (0.0185) |
| Protan | 0.0754 (0.0593) | 0.1414 (0.0276) | 0.1418 (0.0269) | 0.0236 (0.0104) | 0.0265 (0.0071) | 0.0259 (0.0018) | 0.0565 (0.0161) | 0.0563 (0.0106) | 0.0525 (0.0121) |
| Deutan | 0.0220 (0.0066) | 0.0213 (0.0045) | 0.0224 (0.0041) | 0.0822 (0.0475) | 0.1097 (0.0411) | 0.1112 (0.0387) | 0.0343 (0.0155) | 0.0469 (0.0142) | 0.0477 (0.0156) |
| **Noise** | L | | | M | | | S | | |
| #of Charts | 1 | 2 | 3 | 1 | 2 | 3 | 1 | 2 | 3 |
| CN | 2.6705 (1.4328) | 2.3261 (1.4703) | 2.6420 (2.3455) | 2.9072 (2.3990) | 2.3711 (2.4799) | 2.7906 (2.8822) | 4.3502 (2.2982) | 5.1306 (4.3419) | 4.9635 (3.5380) |
| Protan | 27.1211 (20.2595) | 25.3432 (22.1363) | 25.7238 (21.3682) | 2.2436 (1.8449) | 3.4263 (1.9815) | 2.2355 (1.5504) | 5.5647 (2.4101) | 5.1143 (4.9758) | 10.7581 (10.6268) |
| Deutan | 4.9900 (1.9075) | 8.2631 (5.1028) | 7.3670 (2.9478) | 10.5348 (13.4108) | 10.0775 (12.7392) | 15.2871 (14.2381) | 4.2203 (3.0147) | 3.6451 (2.0066) | 4.3365 (3.6860) |
| **Slope** | L | | | M | | | S | | |
| #of Charts | 1 | 2 | 3 | 1 | 2 | 3 | 1 | 2 | 3 |
| CN | 0.0146 (0.0098) | 0.0122 (0.0055) | 0.0115 (0.0042) | 0.0147 (0.0091) | 0.0120 (0.0040) | 0.0121 (0.0059) | 0.0332 (0.0398) | 0.0244 (0.0259) | 0.0236 (0.0195) |
| Protan | 0.0100 (0.0000) | 0.0100 (0.0000) | 0.0117 (0.0041) | 0.0113 (0.0031) | 0.0111 (0.0026) | 0.0118 (0.0043) | 0.0345 (0.0505) | 0.0151 (0.0067) | 0.0170 (0.0078) |
| Deutan | 0.0192 (0.0130) | 0.0121 (0.0064) | 0.0109 (0.0026) | 0.0173 (0.0116) | 0.0129 (0.0044) | 0.0112 (0.0024) | 0.0361 (0.0356) | 0.0251 (0.0219) | 0.0226 (0.0185) |

*Table S6: AIM Color Detection trial-by-trial data for all three metrics (threshold, noise, and slope) in mean(std).*

| **Threshold** | Red | | | RP | | |
|---|---|---|---|---|---|---|
| #of Charts | 1 | 2 | 3 | 1 | 2 | 3 |
| CN | 0.4640 (0.2248) | 0.5537 (0.1459) | 0.5517 (0.1414) | 0.2045 (0.1242) | 0.2467 (0.1115) | 0.2568 (0.0924) |
| Protan | 0.4780 (0.2641) | 0.5618 (0.1739) | 0.5595 (0.1711) | 0.2305 (0.1344) | 0.2470 (0.1225) | 0.2719 (0.0950) |
| Deutan | 0.5983 (0.2801) | 0.6996 (0.2706) | 0.6994 (0.1997) | 0.2006 (0.0945) | 0.2181 (0.0893) | 0.2580 (0.0461) |
| | Purple | | | PG | | |
| #of Charts | 1 | 2 | 3 | 1 | 2 | 3 |
| CN | 0.0682 (0.0480) | 0.0675 (0.0304) | 0.0700 (0.0247) | 0.1332 (0.0915) | 0.1414 (0.0777) | 0.1258 (0.0664) |
| Protan | 0.2886 (0.1366) | 0.3386 (0.1028) | 0.4112 (0.0539) | 0.1806 (0.0797) | 0.1535 (0.0558) | 0.1559 (0.0462) |
| Deutan | 0.2436 (0.1287) | 0.3828 (0.2309) | 0.4151 (0.2603) | 0.1632 (0.0705) | 0.1818 (0.0399) | 0.1637 (0.0245) |
| | Green | | | GY | | |
| #of Charts | 1 | 2 | 3 | 1 | 2 | 3 |
| CN | 0.4313 (0.1639) | 0.3967 (0.1715) | 0.4622 (0.1324) | 0.1885 (0.1240) | 0.2041 (0.0874) | 0.1957 (0.0799) |
| Protan | 0.5475 (0.3040) | 0.7109 (0.1753) | 0.7505 (0.1763) | 0.1868 (0.1101) | 0.1865 (0.0824) | 0.1542 (0.1138) |
| Deutan | 0.5264 (0.2574) | 0.6067 (0.1685) | 0.6280 (0.1195) | 0.0981 (0.0723) | 0.1633 (0.0682) | 0.1767 (0.0495) |
| | Yellow | | | YR | | |
| #of Charts | 1 | 2 | 3 | 1 | 2 | 3 |
| CN | 0.0468 (0.0297) | 0.0459 (0.0249) | 0.0484 (0.0168) | 0.1914 (0.1064) | 0.2121 (0.0522) | 0.2088 (0.0519) |
| Protan | 0.2985 (0.1720) | 0.3796 (0.1034) | 0.4054 (0.0854) | 0.2401 (0.1411) | 0.2622 (0.0768) | 0.2587 (0.0785) |
| Deutan | 0.2929 (0.2496) | 0.4778 (0.2486) | 0.4531 (0.2518) | 0.1734 (0.1265) | 0.1954 (0.0986) | 0.2103 (0.0812) |

*Table S7: AIM Color Discrimination trial-by-trial data for threshold in mean(std).*

| **Noise** | Red | | | RP | | |
|---|---|---|---|---|---|---|
| #of Charts | 1 | 2 | 3 | 1 | 2 | 3 |
| CN | 12.7681 (6.6419) | 10.7174 (5.1111) | 11.4096 (4.5485) | 7.2752 (3.3657) | 7.1613 (3.4335) | 8.0619 (3.6122) |
| Protan | 8.9976 (2.9260) | 10.1891 (4.4125) | 12.2420 (5.4573) | 7.4132 (4.8024) | 6.8974 (3.2244) | 5.9660 (2.1286) |
| Deutan | 12.2612 (6.4681) | 8.4397 (4.0082) | 10.3167 (5.0798) | 7.6288 (4.3173) | 8.4062 (4.0942) | 6.5966 (1.9352) |
| | Purple | | | PG | | |

| #of Charts | 1 | 2 | 3 | 1 | 2 | 3 |
|---|---|---|---|---|---|---|
| CN | 4.5507 (2.7432) | 4.8807 (2.9049) | 4.5854 (2.2885) | 8.7528 (6.5441) | 5.2438 (3.0446) | 7.0564 (5.4705) |
| Protan | 16.1756 (7.1774) | 9.9544 (7.7347) | 8.1042 (7.8756) | 10.5572 (7.1343) | 6.0641 (3.4690) | 6.7812 (2.1931) |
| Deutan | 19.6926 (4.7218) | 13.9087 (7.1197) | 12.8628 (4.7685) | 11.6032 (8.2568) | 5.3371 (2.1820) | 5.3610 (2.4609) |
| | Green | | | GY | | |
| #of Charts | 1 | 2 | 3 | 1 | 2 | 3 |
| CN | 8.8324 (4.3207) | 10.3999 (4.8067) | 9.2764 (3.3414) | 6.8366 (6.1153) | 6.5833 (3.8292) | 7.1592 (5.0860) |
| Protan | 10.4346 (3.8601) | 9.2136 (1.8901) | 9.2109 (2.6524) | 8.6676 (6.6338) | 6.4806 (2.3792) | 8.6037 (6.1278) |
| Deutan | 8.9143 (6.1254) | 7.0270 (3.6216) | 6.9554 (3.0493) | 7.9977 (4.4278) | 6.8022 (1.8919) | 7.2234 (2.0778) |
| | Yellow | | | YR | | |
| #of Charts | 1 | 2 | 3 | 1 | 2 | 3 |
| CN | 4.6633 (3.0666) | 4.9028 (2.6249) | 4.9974 (2.4312) | 5.9210 (2.5778) | 6.2056 (2.6551) | 6.7775 (2.5385) |
| Protan | 18.7536 (5.8059) | 11.3254 (7.7378) | 8.1277 (3.9774) | 7.2426 (2.6523) | 6.7575 (2.2741) | 6.4897 (2.1307) |
| Deutan | 19.3326 (7.1669) | 19.1403 (6.8417) | 19.1297 (3.4657) | 5.5011 (2.0579) | 6.2630 (1.9214) | 6.7001 (3.3838) |

*Table S8: AIM Color Discrimination trial-by-trial data for noise in mean(std).*

| **Slope** | Red | | | RP | | |
|---|---|---|---|---|---|---|
| #of Charts | 1 | 2 | 3 | 1 | 2 | 3 |
| CN | 0.0596 (0.0525) | 0.0871 (0.0595) | 0.0923 (0.0573) | 0.0802 (0.0617) | 0.0723 (0.0567) | 0.0707 (0.0457) |
| Protan | 0.0805 (0.0539) | 0.0700 (0.0567) | 0.1002 (0.0603) | 0.1014 (0.0562) | 0.0836 (0.0539) | 0.1103 (0.0492) |
| Deutan | 0.0724 (0.0550) | 0.0638 (0.0567) | 0.0924 (0.0511) | 0.0548 (0.0564) | 0.0407 (0.0427) | 0.0907 (0.0485) |
| | Purple | | | PG | | |
| #of Charts | 1 | 2 | 3 | 1 | 2 | 3 |
| CN | 0.0478 (0.0441) | 0.0357 (0.0305) | 0.0387 (0.0310) | 0.0550 (0.0465) | 0.0579 (0.0456) | 0.0527 (0.0431) |
| Protan | 0.0586 (0.0593) | 0.0772 (0.0498) | 0.0860 (0.0550) | 0.0850 (0.0712) | 0.1064 (0.0669) | 0.0875 (0.0685) |
| Deutan | 0.0608 (0.0574) | 0.0803 (0.0608) | 0.1174 (0.0480) | 0.0489 (0.0573) | 0.0537 (0.0467) | 0.0705 (0.0574) |
| | Green | | | GY | | |
| #of Charts | 1 | 2 | 3 | 1 | 2 | 3 |
| CN | 0.0575 (0.0562) | 0.0706 (0.0605) | 0.0910 (0.0566) | 0.0775 (0.0602) | 0.0895 (0.0574) | 0.0829 (0.0537) |
| Protan | 0.0702 (0.0580) | 0.0489 (0.0509) | 0.1039 (0.0435) | 0.0747 (0.0640) | 0.1271 (0.0355) | 0.0789 (0.0573) |

| | | | | | | |
|---|---|---|---|---|---|---|
| Deutan | 0.0601 (0.0536) | 0.0751 (0.0619) | 0.0680 (0.0596) | 0.0633 (0.0650) | 0.0721 (0.0565) | 0.0602 (0.0498) |
| | Yellow | | | YR | | |
| #of Charts | 1 | 2 | 3 | 1 | 2 | 3 |
| CN | 0.0382 (0.0319) | 0.0301 (0.0187) | 0.0243 (0.0106) | 0.0756 (0.0606) | 0.0782 (0.0579) | 0.0768 (0.0505) |
| Protan | 0.0851 (0.0585) | 0.1027 (0.0500) | 0.1237 (0.0520) | 0.0591 (0.0590) | 0.0815 (0.0512) | 0.0829 (0.0481) |
| Deutan | 0.0890 (0.0661) | 0.0574 (0.0492) | 0.0763 (0.0589) | 0.0873 (0.0615) | 0.0875 (0.0533) | 0.0779 (0.0566) |

*Table S9: AIM Color Discrimination trial-by-trial data for slope in mean(std).*
